# An overlapped natural antisense lncRNA FCER1A-AS controls anaphylactic reaction by promoting FcεRIα expression

**DOI:** 10.1101/2022.04.05.487139

**Authors:** Ruo-yu Tang, Lan Yin, Liang Yao, Xiao-Ping Chen

**Author notes:** **Corresponding author:** Xiao-Ping Chen, MD, PhD, Professor.

## Abstract

Functioned as α-subunit of the high-affinity immunoglobulin E receptor (Fcε RIα), Fcε RIα plays a central role in the pathogenesis of Ig-E-mediated allergy and other IgE-related disorders. Fcε RIα is normally expressed only in limited spectrum of cells like basophils and mast cells, but the mechanism of controlling Fcε RIα expression in these cells is less well understood. In this study, we found fully overlapped natural antisense transcript (NATs) of Fcε RIα (*FCER1A*-AS) is co-expressed with cognate sense transcript (*FCER1A*-S) in IL-3 induced Fcε RIα-expressing cells or in high Fcε RIα-expressing cell line MC/9. When *FCER1A*-AS is selectively knocked down by CRISPR/RfxCas13d (CasRx) approach in MC/9, expression of mRNA and proteins of *FCER1A*-S is also markedly decreased. Furthermore, deficiency of *FCER1A*-AS along with lack of *FCER1A*-S expression is found in two different lines of gene-targeted mice in which *FCER1A* locus is disturbed at different sites. More importantly, the *FCER1A*-AS-deficient homozygous mice display similarly diminished anaphylactic reaction as *FCER1A* gene knockout mice. Thus, we uncovered in this investigation that the expression of *FCER1A*-S is positively regulated by co-expressed fully overlapped antisense transcript.

## Introduction

The high affinity receptor for immunoglobulin E (FcεRI) is known to be responsible for IgE-dependent immunity, IgE-mediated hypersensitivity and probably for some antibody-mediated autoimmune diseases (AAID) (Charles, 2021). Activation of FcεRI triggers degranulation releasing granule contents containing early and late inflammatory mediators like histamines, arachidonic acid derivatives and cytokines. Therefore, block of FcεRI activation by anti-IgE monoclonal antibodies are proved to be clinically effective in treatment of atopic diseases such as asthma, atopic dermatitis, chronic urticaria, and even AAID (Okayama et al., 2020)

Each FcεRI is a tetrameric molecule composed of one α, one β chain and two γ chains[αβγ2](Blank et al., 1989). The chain (FcεRIα) confers the receptor with high affinity binding with Fc portion of IgE; β chain amplifies the signal and γ chains mediate signal transduction. FcεRIα is almost exclusively expressed on mast cells and basophils in both mouse and human. Recent studies, however, suggested that FcεRIα may also be a marker for type 2 immune response, since its expression is also found on type 2 inflammation-associated dendritic cells(Hammad et al., 2010; Ma et al., 2015) or monocytes(Satoh et al., 2017).

IL-3-GATA-2 axis appears to dictate the limited expression spectrum of FcεRIα on lineages of mast cells and basophils (Li, Qi, Liu, & Huang, 2015; Ohmori et al., 2015). But the mechanisms that how FcεRIα expression is controlled in these cells are unknown. Human and mouse *FCER1A* gene transcripts are composed of 5 exons along with 5’ UTR and 3’ UTR. Earlier on in our attempts to investigate the role of FcεRIα-positive cells in vivo, we constructed FcεRIα-DTR mice (knock-in DTR gene at 3’UTR of *FCER1A*) in hope that FcεRIα-expressing cells are specifically abrogated upon administration of DT. Unexpectedly, these FcεRIα-DTR homozygous mice are defective of FcεRIα expression on cell surfaces due to failure of *FCER1A* transcription. Another attempt was pursued to replace the whole mouse *FCER1A* gene with DTR gene. Failure of transcription of edited region was again observed. In both cases, lineage differentiation of FcεRIα-expressing cells is not affected. Thus, in addition to trans-regulation by GATA-2, mouse *FCER1A* expression may also be regulated by unidentified sequence-dependent cis-regulation.

In this study we found that when mouse *FCER1A* gene is transcribed, a fully complementary noncoding *FCER1A* antisense RNA (*FCER1A*-AS) is also transcribed simultaneously. The gene editing procedures performed disrupted the transcription of *FCER1A*-AS. Further investigations showed that these natural full-length antisense transcripts play positive role in regulation of sense *FCER1A* expression. Our results provide a novel regulation pathway for allergy-controlling molecule FcεRIα. In addition, our data also suggested that cautions need to be taken while gene editing is executed on an allele which expression is subjected to cis regulation by antisense RNA.

## Result

### Identification of mouse *FCER1A* natural antisense transcript

In order to investigate the regulation of *FCER1A* mRNA expression, we obtained FcεRIα-positive cells from IL-3-treated bone marrow cultures to overcome low levels of these cells under steady state. Presence of IL-3 markedly increased the percentage of FcεRIα-positive cells 6 days after treatment, whereas GM-CSF did not induce any FcεRIα-positive cells which was used as experimental negative control (Fig1-supplementary 1). Primers specific for sense strand or putative antisense strand were designed for strand-specific reverse transcription. PCR amplification of strand-specific cDNAs was then performed with multiple sets of primers covering different exons or UTR regions either on sense strand or on putative antisense strand (Fig.1a). *FCER1A* sense transcripts(*FCER1A*-S)were almost undetectable in bone marrow cells and were readily induced in the presence of IL-3 as expected (Fig. 1b and 1d). Antisense transcripts of *FCER1A*(*FCER1A*-AS)which were barely detectable in freshly isolated bone marrow cells were also induced with similar time kinetics as *FCER1A*-S following IL-3 treatment (Fig. 1c and 1d). GM-CSF treatment induced neither sense nor antisense transcripts of *FCER1A* (Fig.1b, 1c and 1d). Presence of FCER1A-AS is also verified on MC/9, a mouse mast cell line which expresses high levels of *FCER1A*-S (Fig1-supplementary 2). Therefore, antisense transcript is expressed in parallel with sense transcript at *FACER1A* locus. When RNA stability of both sense and antisense was examined in MC/9 cells, half-life for *FCER1A*-AS was approximate 13 h, nearly 3 h longer than *FCER1A*-S (Fig1-supplementary 3).

**Figure 1.**
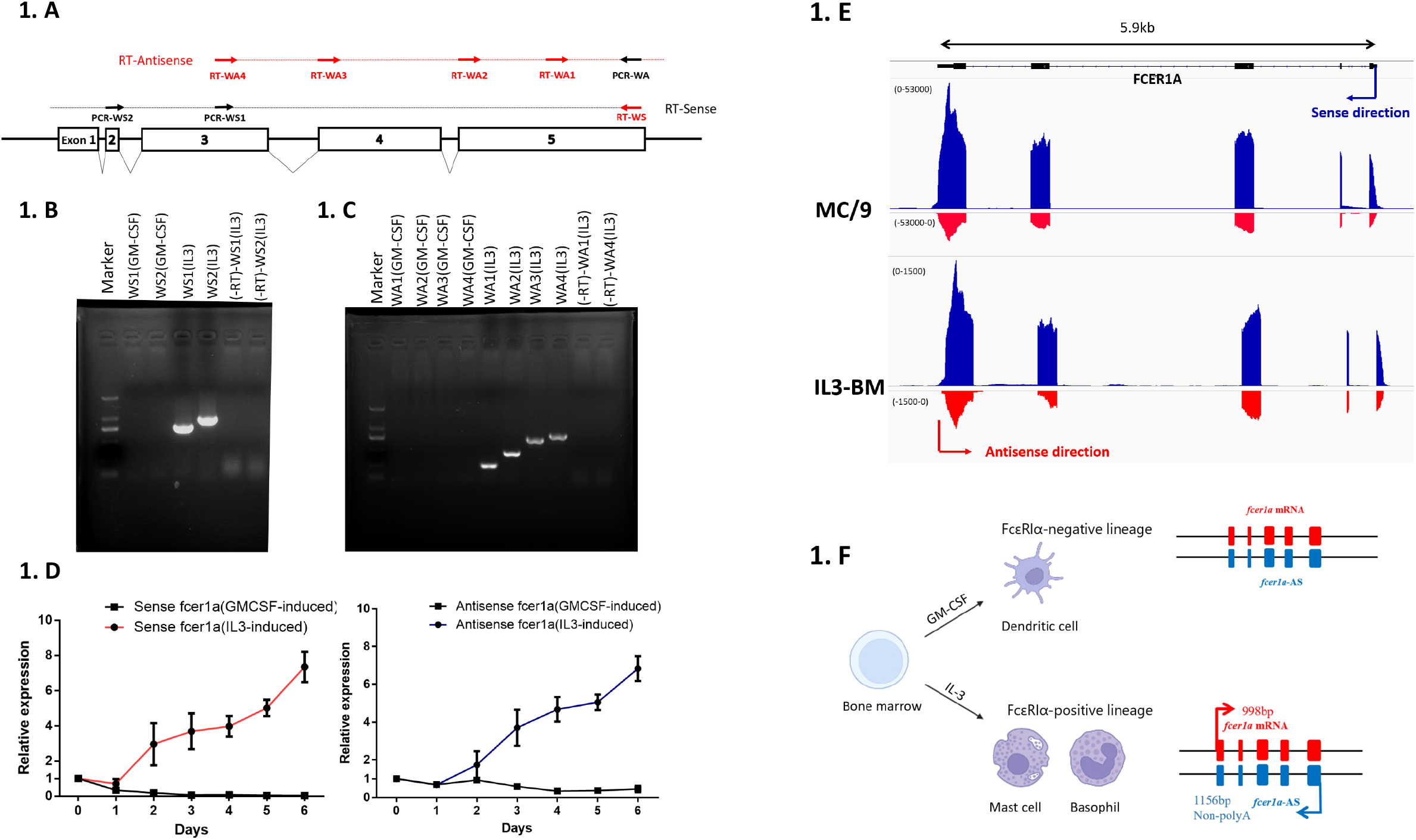
Fully complementary antisense transcript of *FCER1A* (*FCER1A*-AS) is expressed along with its sense transcript. **(A)** RT-PCR primers design for detection of sense (Black dotted-line) and antisense (red dotted-line) of *FCER1A*. RT primers (red arrows) for sense (WS) or antisense (WA1 to WA4) and PCR primers (black arrows). **(B and C)** Gel analysis of RT-PCR products of sense or antisense of *FCER1A* from 6 days of IL-3- or GM-CSF-treated bone marrow cells. The -RT column shows PCR amplifications of DNase-treated RNA without RT for genomic DNA contamination control. **(D)** Time kinetics of sense (left panel) and antisense (right panel) of *FCER1A* expression from 6 days of IL-3 (circle) - or GM-CSF (square)-treated bone marrow cells. Combined data shows relative expression in relation to expression level at time zero. n=4 independent animals. **(E)** Strand-specific RNA-seq profiles of *FCER1A* genes from MC/9 and IL-3-treated bone marrow cells. Y axis reflects counts of RNA-seq reads. Arrows indicate transcription direction of sense (blue) or antisense (red). Top black lines depict known *FCER1A* gene reference. **(F)** Schematic illustration of *FCER1A*-S and *FCER1A*-AS being co-expressed only in IL-3 induced FcεRIα-postive cells. *FCER1A-AS* is 158bp longer than its sense counterpart without polyA tail

Rapid amplification of cDNA ends (RACE) was performed on RNA isolated from MC/9 cells or from IL-3-induced bone marrow cells to identify the starting and ending site of *FCER1A*-AS transcription. Results demonstrated that *FCER1A*-AS is 158bp longer than its sense counterpart without polyA tail (Fig1-supplementary 4). Sanger sequencing of PCR products of *FCER1A*-AS revealed that the antisense transcript fully overlaps with sense transcript including all five exons together with 5’ untranslated region (UTR) and 3’UTR (Fig1-supplementary 4). The existence of *FCER1A*-AS and its fully overlapping feature with *FCER1A*-S were both demonstrated by strand-specific RNA sequencing by next generation sequencing (NGS) platform together with bioinformatics analysis (Fig.1e). There is no coding capacity predicted on *FCER1A*-AS in silico (Fig1-supplementary 5, http://cpc2.gao-lab.org/)(Kang et al., 2017). These data confirmed that a fully overlapped, non-coding, natural antisense transcript of Fcε RIα (*FCER1A*-AS) is co-expressed with cognate sense transcript (*FCER1A*-S) in IL-3 induced Fcε RIα-expressing cells (Fig.1f).

### Knockdown of *FCER1A*-AS reduces FcεRIα mRNA and protein expression in vitro

To explore if *FCER1A*-AS plays any regulatory roles in expression of *FCER1A*-S, we designed RNA knockdown on *FCER1A*-AS in MC/9 cells. Considering the full-length overlapping feature between *FCER1A*-AS and *FCER1A*-S and unknowns on *FCER1A*-AS RNA structures, we employed recently developed CRISPR/RfxCas13d (CasRx) technique to selectively knock down of antisense strand (He et al., 2020; Konermann et al., 2018; Zhou et al., 2020). Four different guide RNAs targeting at 5’ regions of *FCER1A*-AS (*FCER1A*-AS-gRNAs) were designed (Fig 2a). Two non-targeting gRNA (NT-gRNAs) that do not target at any known sequences in mouse genomes were used as controls (Fig 2a). Plasmids expressing gRNAs or CasRx were co-transfected into MC/9 cells by electroporation. Cells transfected with *FCER1A*-AS-gRNAs or NT-gRNAs displayed similarly affected viability (Figure2-supplementary 1). Significant knockdown of *FCER1A*-AS by all four *FCER1A*-AS-gRNAs but not by NT-gRNAs were all observed 48 hrs after co-transfection (Figure2-supplementary 2). Unexpectedly however, significantly reduced positivity of FcεRIα-expressing MC/9 cells, markedly decreased levels of FcεRIα expression on cell surfaces measured by mean fluorescence intensity and of FcεRIα transcription were also found in successfully co-transfected cells in which *FCER1A*-AS was the molecule being knocked down (Fig. 2b, 2c, 2d, Figure2-supplementary 3). No inhibition on expression of sense transcript was found when NT-gRNAs/CasRx was transfected. No further inhibition was seen when multiple sites on *FCER1A*-AS were targeted by multiple gRNAs strings. Moreover, knock down of *FCER1A*-AS did not result alterations on other known mast cell-associated molecules examined including GATA-2 and CD200R3 (Fig, 2f and 2e). Therefore, *FCER1A*-AS plays a critical positive role in regulation of the transcription of its opposite sense strand.

**Figure 2.**
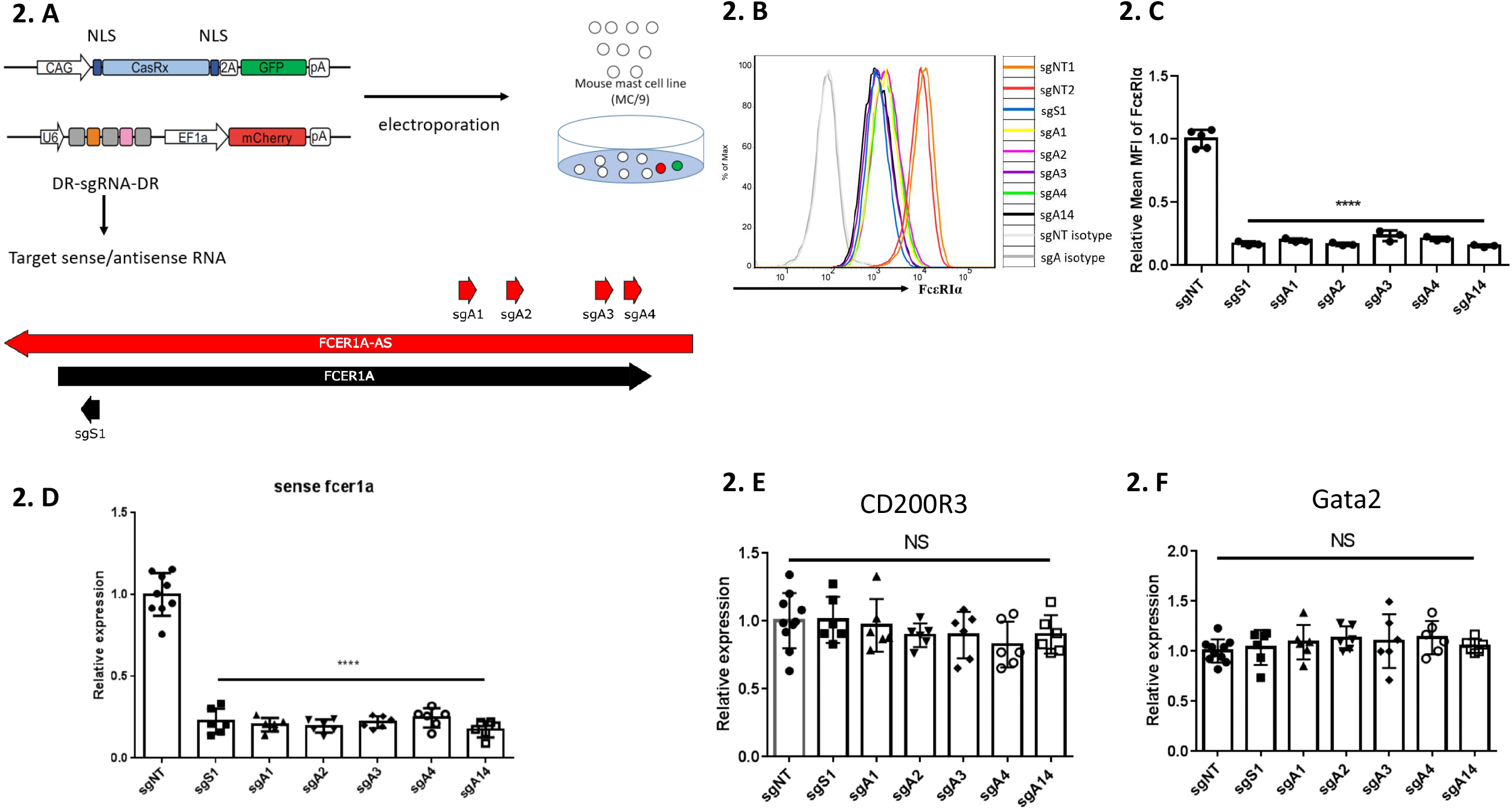
CasRx-mediated knockdown of *FCER1A-S*/*FCER1A*-AS in MC/9. **(A)** Strategies for CasRx-mediated knockdown of *FCER1A*-S or *FCER1A*-AS in MC/9. Four differently targeted sgRNAs (A1, A2, A3, and A4) and one multiplexed sgRNA combining all 4 SgRNAs (A1-4) were designed for knockdown of *FCER1A*-AS, and one sgRNA (S1) for *FCER1A*-S. MC/9 cells were co-transfected by electroporation with expression plasmids for sgRNA and CasRx. **(B)** FACS analysis of surface expression of FcεRIα following *FCER1A*-AS-sgRNA (sgA) or FCER1A-S-sgRNAs (sgS) transfection. Shown are FACS staining of co-transfected cells gated on cells positive both for GFP+ and Mcherry+ with non-targeting sgRNAs (sgNT) and isotype staining (grey) as controls. Histogram shown is representative of three independent experiments. **(C)** Summarized relative mean fluorescence intensity (MFI) of FACS staining of FcεRIα over MFI of sgNT. n=3 independent experiments. ****P < 0.0001. **(D)** Strand-specific quantitative RT-PCR of *FCER1A*-S expression following *FCER1A*-AS-sgRNA (sgA) or FCER1A-S-sgRNAs (sgS) transfection. Relative expression indicates the reduced qRT-PCR levels in each sgRNA knock down treatment relating to control sgNT. Data are mean ± SD of six different experiments. ****P < 0.0001. **(E and F)** Quantitative RT-PCR of *GATA2* and *CD200R3* in MC/9 cells following *FCER1A*-AS-sgRNA (sgA) or FCER1A-S-sgRNAs (sgS) transfection. RNA from each group was reverse transcribed by random hexamers. Shown is the expression levels relative to control sgNT. n=6 different experiments. NS: no significant differences

### Deficiency of *FCER1A*-AS results loss of *FCER1A*-S in vivo

Two different gene-editing mice targeted at *FCER1A* locus constructed by us were found to be lack of antisense transcript expression. One line was constructed by incorporating sequences encoding human DTR and self-cleaving peptide (P2A) right before 3’ UTR without disturbing coding sequences of *FCER1A* gene (Fig. 3a, FCER1A^dtr/dtr^ mice, Model I). The other line was to replace whole *FCER1A* locus with DTR-P2A-GFP module with UTR regions undisturbed (Fig. 3b, FCER1A^dtrGFP/dtrGFP^, Model II). Mice of both lines are fertile and healthy. The genotyping of both lines were depicted (Fig. 3c and 3d) and edited genetic region were confirmed by sequencing. We initially built these 2 lines in order to investigate the role of FcεRIα-positive cells in vivo, in hope that FcεRIα-expressing cells are specifically abrogated upon administration of DT. However, we found that the expression of antisense transcripts was deficient in homozygous mice from both models using primers designed specific for strand-specific reverse transcription in either lines (Fig 4a and 4b).

**Figure3.**
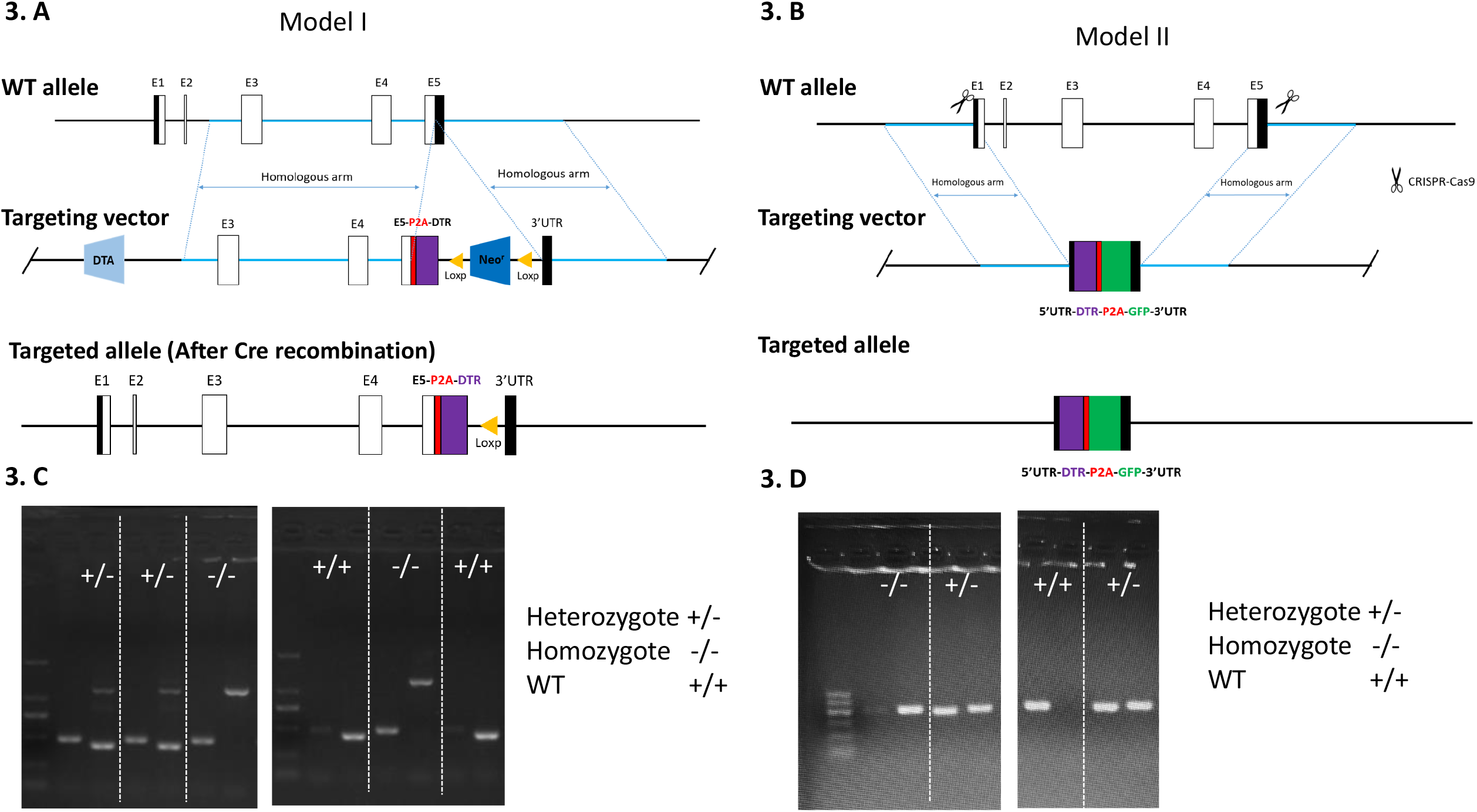
Generation of two gene-editing mouse models at *FCER1A* locus and their genotyping. **(A)** Targeting strategy for FCER1A^dtr/dtr^ (model I). FCER1A^dtr/dtr^ mice were generated via introducing a P2A-DTR-loxP-Neor-loxP cassette into 3’UTR of FCERIA allele (before stop codon) by homologous recombination. The coding sequence remained intact. **(B)** Targeting strategy for FCER1A^dtrGFP/dtrGFP^ (model II). FCER1A^dtrGFP/dtrGFP^ mice were generated via replacing the whole FCER1A region with DTR-P2A-GFP module without disruption of 5’UTR or 3’UTR by CRISPR-Cas9 strategy. **(C and D)** Genotyping of FCER1A^dtr/dtr^(model I) andFCER1A^dtrGFP/dtrGFP^ (model II) mice as depicted. For model I, PCR was carried out using primer pairs Region2-F/Region2-R and Region2-F/Region1-R. For model II, PCR was carried out using primer pairs DTR2AGFP-F/DTR2AGFP-R and FCER1Ain-F/ FCER1Ain-R

**Figure 4.**
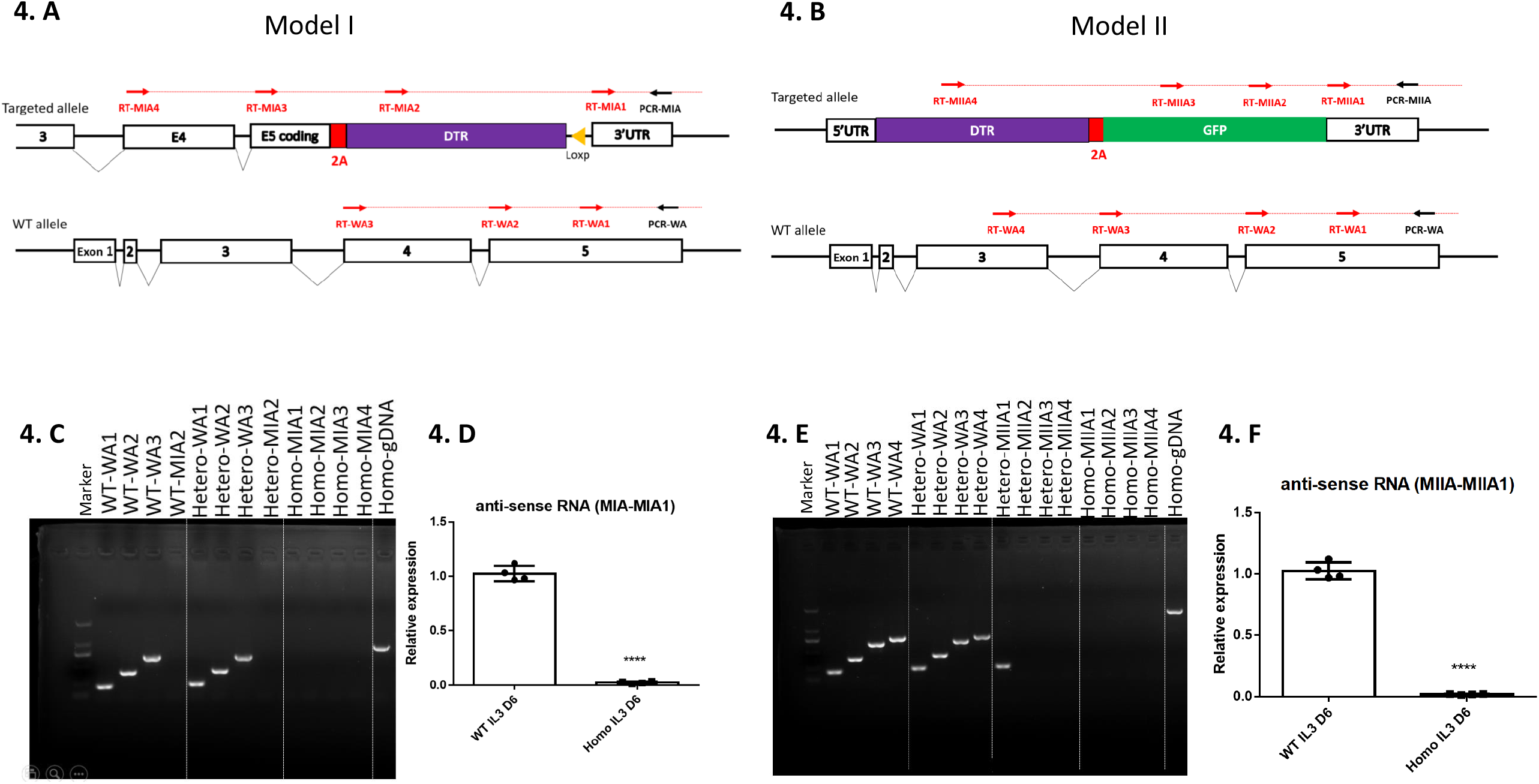
Deficiency of *FCER1A*-AS in two *FCER1A* gene editing mouse models. **(A and B)** RT-PCR primers design for detection of antisense transcripts at *FCER1A* loci in WT, heterozygous and homozygous mice of Model I (A) and Model II (B). RT primers (red arrows) in WT (WA1 to WA4), Model I (MIA1 to MIA4) and Model II (MIIA1 to MIIA4). Common PCR primer (black arrows) is used together with respective RT primers to amplify RT products. **(C to F)** RT-PCR of antisense transcripts of *FCER1A* loci from 6 days of IL3 -treated bone marrow cells in WT, Model I and Model II. Gel analysis of RT-PCR products of antisense transcripts of FCER1A loci in **(C)** WT (WA1 to WA3, MIA2), Model I heterozygous (WA1 to WA3, MIA2), Model I homozygous (MIA1 to MIA4) and **(E)** WT (WA1 to WA4), Model II heterozygous (WA1 to WA4, MIIA1 to MIIA4), Model II homozygous (MIIA1 to MIIA4). Genomic DNA (gDNA) of homozygous mice was included (Model I: MIA2-MIA; Model II: MIIA4-MIIA) for positive control of PCR reaction. Strand-specific quantitative RT-PCR of antisense transcripts of *FCER1A* loci from 6 days of IL3-treated bone marrow cells in WT, Model I (**D)** and Model II (**F)**. RNA from each group was reverse transcribed by common antisense-specific primer (RT-MIA1/MIIA1), and the region of 3’UTR was selected for final qPCR detection. Relative expression indicates the ratio relative to WT. Data are the mean ± SD of four separate animals from each model. ****P < 0.0001.

Significantly, the expression of sense transcript is also affected in both models. The cell surface expression of FcεRIα was markedly decreased in FCER1A^dtr/dtr^ Model I homozygous animals when FcεRIα expression was examined in peripheral blood, in IL-3-treated bone marrow cultures, or in helminth infected mice in which FcεRIα-positive cells are known to be elevated(Ma et al., 2015)(Fig. 5a). Transcription of sense transcripts, as shown in Fig. 5b, was almost diminished in homozygous Model I animals, whereas 50% reduction of sense transcript expression was seen in heterozygous mice (Fig. 5b). Additionally, inserted gene DTR was not able to be transcribed in this model (Fig. 5b). When the expression of inserted elements, *ie*, DTR or GFP, was assessed in Model II, they were not detectable in homozygous or in heterozygous mice (Fig. 5c and 5d). In both cases, lack of expression of sense transcription is not due to lack of IL-3 responsiveness, because the number of basophils measured by CD200R3 and CD49b in peripheral blood or expression of CD200R3 in IL-3-treated bone marrow cells were not affected (Figure5-supplementary 1).

**Figure 5.**
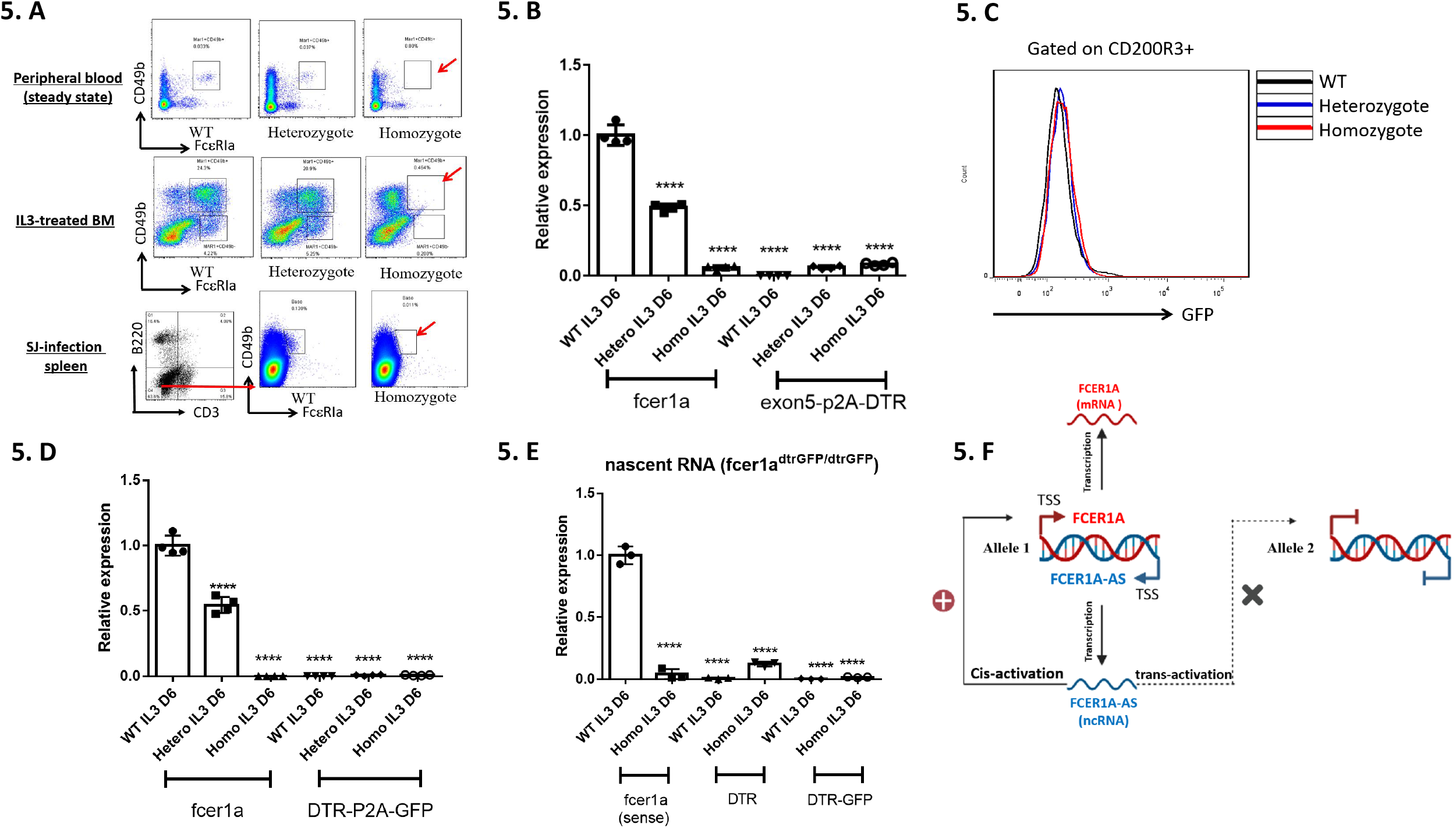
Absence of *FCER1A*-AS is associated with loss of *FCER1A*-S expression and anaphylactic reaction in vivo. **(A)** FACS analysis of cell surface expression of FcεRIα in normal peripheral blood cells, IL3-induced bone marrow cells and *Schistosoma japonicum* (Sj) infected splenocytes in mice from WT, heterozygous Model I and homozygous Model I. In case of Sj infection, detection of FcεRIα-positive cells was gated on NBNT (CD3-B220-) cells. **(B)** Quantitative RT-PCR of *FCER1A*-S and inserted DTR from 6 days of IL3-treated bone marrow cells in WT, heterozygous Model I and homozygous Model I. RNA was reverse transcribed by oligo dT. Shown is expression level in each setting relative to *FCER1A*-S level at day 6 in WT. Data are mean ± SD of four separate animals from each model. ****P < 0.0001. **(C)** FACS examination of cells expressing inserted GFP in IL3-induced bone marrow cells from mice of WT, heterozygous Model II and homozygous Model II. Shown are cells gated on CD200R3 positive populations. **(D)** Quantitative RT-PCR analysis of *FCER1A*-S and replaced DTR-GFP fusion transcripts from 6 days of IL3-treated bone marrow cells in WT, heterozygous Model II and homozygous Model II. RNA from each groups was reverse transcribed by oligo dT. After GAPDH normalization, relative expression is calculated to the expression of *FCER1A*-S at day 6 in WT. Data are the mean ± SD of four separate animals from each model. ****P < 0.0001. **(E)** Quantitative RT-PCR of nascent transcripts of *FCER1A* or inserted DTR and DTR-GFP from 6 days of IL3-treated bone marrow cells in WT and homozygous Model II. RNA from each group was reverse transcribed by random hexamers. Relative expression is calculated to the expression of relevant molecules at day 6 in WT. Data are the mean ± SD of four separate animal s from each model. ****P < 0.0001. **(F)** Schematic depiction of critical positive role of FCER1A-AS in regulation of the transcription of FCER1A-S via cis-activation.

We further performed click-iT nascent RNA capture assay in IL-3-induced bone marrow cells from Model II mice to explore if defect in sense transcript transcription is caused by failure of transcription initiation. As shown in Fig. 5e, nascent RNA was readily detectable in cells from wild type mice, indicating IL-3 treatment promotes sense gene transcription as expected. When similarly treated bone marrow cells of homozygous model II mice were studied, the levels of nascent RNA for two inserted gene transcripts, ie, DTR or DTR-GFP, were much lower than the levels of nascent RNA found in unedited wild type animals (Fig. 5e).

Taken together, we believed that both gene targeting procedures disrupt the antisense transcription hence failure of sense transcription initiation. It is noteworthy to mention that transcripts of antisense were still detectable in heterozygous mice (Fig. 4c, 4d, 4e and 4f), indicating the disrupted machinery by gene editing procedures required for antisense transcription only acts on affected chromosome. Furthermore, the existence of FCER1A-AS from sister chromosome is unable to rescue the inhibited transcription of sense transcripts on the other chromosome in heterozygote of both lines (Fig 4c, 4e, 5a, 5b, 5c, 5d). Therefore, positive regulation of S by AS appears to act in a cis-acting manner. These findings together indicated that sequence intrinsic mechanisms may exist to determine the transcription of antisense transcripts at *FCER1A* locus which is required for transcriptional initiation of *FCER1A*-S via cis-acting manner in vivo (Fig. 5f).

### Loss of *FCER1A*-AS markedly reduces anaphylactic responses in vivo

To investigate if loss of *FCER1A*-AS will lead to deficiency in allergic responses, well-established FcεRIα-mediated passive cutaneous anaphylaxis (PCA) reaction was studied in both models. Sensitization by subcutaneous injection of anti-DNP IgE at ears followed by an intravenous administration of antigen DNP is known to induce markedly extravasation at sensitized ears. The severity of PCA is reflected by the level of Evans blue extravasation at ears, the dye given together with DNP via blood. As expected, knockout of *FCER1A* derived from homozygous model II mice led to a 10 fold decrease of extravasation by visualization or by quantification of the amount of Evans blue at sensitized ears (Fig. 6a and 6b). More significantly, equally diminished extravasation was also found in PCA reaction in FCER1A^dtr/dtr^ (Model I) homozygous mice (Fig. 6a and 6b). Since the coding sequences for Fcε RIα in FCER1A^dtr/dtr^ remain intact, loss of PCA responses in this model strongly indicated a critical role played by *FCER1A*-AS in vivo. In addition, we also examined if loss of *FCER1A*-AS affects anti-helminth immunity in which FcεRIα-mediated mechanisms are proposed (Joseph et al., 1997). After Schistosome *japonica*(Sj)infection, FCER1A^dtr/dtr^ model I mice displayed slightly reduced survival comparing to wild type counterparts (Figure6-supplementary 1). Taken together, transcription of *FCER1A*-AS, just like its counterpart *FCER1A*-S, is required for the occurrence of IgE-mediated anaphylactic reactions and anti-helminth immunity.

**Figure 6.**
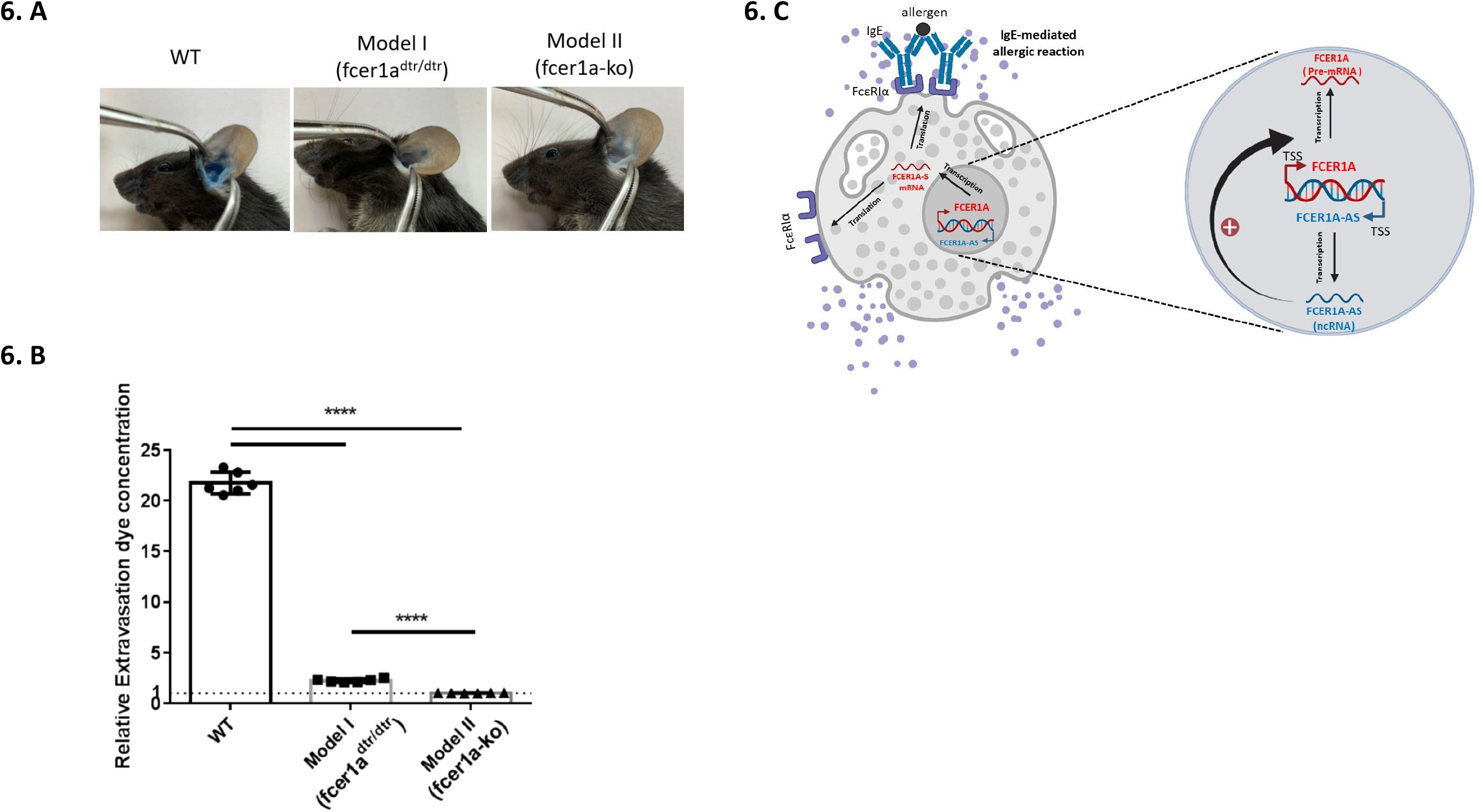
Loss of *FCER1A*-AS markedly reduces anaphylactic responses in vivo. **(A)** Representative photographs of ear extravasation of dye in different murine models during passive cutaneous anaphylaxis. **(B)** Measurement of Evan’s blue extravasation into the ears of mice among WT, Model I homozygous and Model II homozygous mice after challenge with antigen. Relative extravasation dye concentration is a ratio relative to the OD650 readings in Model II homozygous mice (equals to *FCER1A* knock out). Data are the mean ± SD from six ears of three mice. ****P < 0.0001. (C) Model depicting the mode of allergy-controlling molecule Fcε RIα is strickly positively controlled by presence of FCERIA-AS.

### Analysis for initiation potentials of FACERI-AS transcription

To identify potential machineries involved in allowing transcription of antisense of FCER1A, we analyzed gene configuration of FCER1A locus in FcεRIα-expressing bone marrow derived mast cells using NCBI Gene Expression Omnibus (GEO) datasets (GSE145542 and GSE145544)(Li et al., 2021). We found that binding signals for H3K4me3 and H3K27ac, two histone modification fashions with upregulating activities on gene transcription, are enriched at not only at 5’promoter regions of *FCER1A-*S as expected but also at putative 5’promoter flanking region of *FCER1A-* AS in BMMC with comparable peak levels (Fig 7a). Similar signal pattern is also found on ATAC-Seq and on CHIP-Seq of GATA-2, the key positive transcriptional factor for *FCER1A-*S and differentiation of FcεRIα-expressing cells (Li et al., 2015; Ohmori et al., 2015). Since the nearest downstream gene OLFR1404 was not expressed in mast cells (Figure7-supplementary 1), we believed that the gene configuration of upstream of *FCER1A-*AS is in favorable status for its transcription.

**Figure 7.**
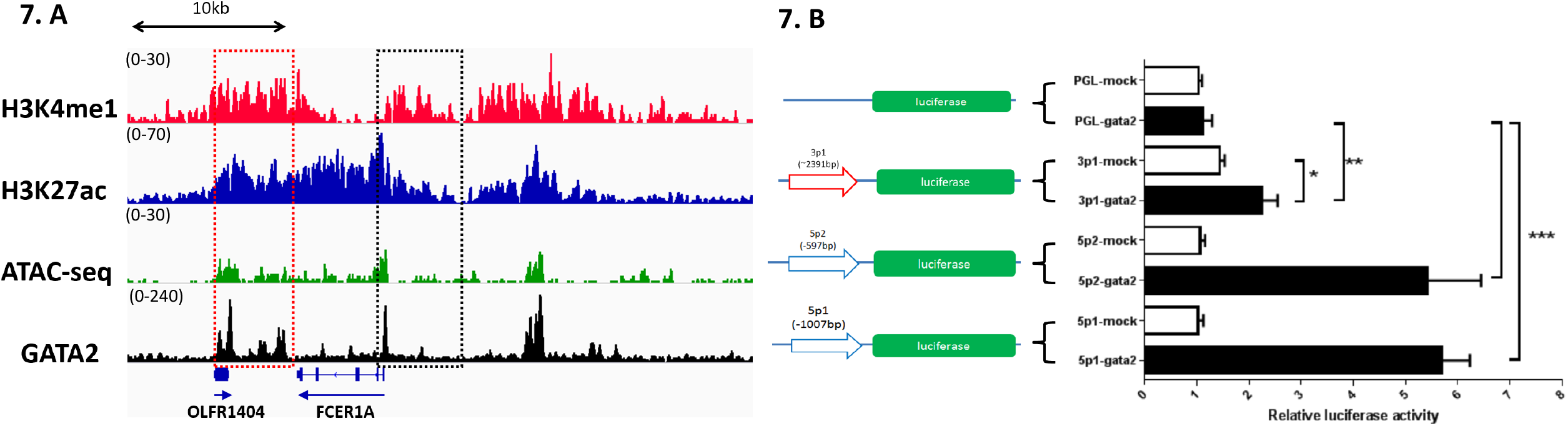
Putative transcription initiation analysis for FACERIA-AS. (A) Representative tracks of H3k4me1/H3k27ac/GATA-2 ChIP-seq and ATAC-seq for FCER1A loucs in bone marrow-derived mast cells. Data are shown as reads density around FCER1A locus. The IGV tracks are generated from one biological sample, representing two biological replicates with similar patterns. The original raw data are retrieved from”GSE145542” and”GSE145544”. (B) Gata-2 transactivates *FCER1A*-AS putative promoter at low level. A reporter plasmid carrying promoter region of the *FCER1A*-S (−1007/+1,5p1; -597/+1,5p2), *FCER1A*-AS (−2391/+1,3p1) or its empty plasmid pGL-4 was transfected into HEK293T cells with GATA-2 expression plasmid pCDNA3.1 (+)-GATA2 (GATA2), pCDNA3.1 (+) (mock). Promoter activities were assessed by luciferase activity compared between GATA-2 and mock. Data are mean ± SD of four independent experiments.

We then examined the potential role of GATA-2 in transactivation of antisense promoter using classic luciferase reporter assay. Weaker GATA-2-associated promoter activities were found 2.4kb downstream of 3’UTR of Fcε RIα gene to allow transcription of *FCERIA*-AS, comparing to significantly presented GATA-2-promoted activities on sense direction (Fig 7b). Therefore, GATA-2 does not appear to function equally at both directions.

## Discussion

In this study we uncovered a novel mechanism of regulation of Fcε RIα by its antisense partner *FCERIA*-AS. Antisense transcripts (AS) or natural antisense transcripts (NATs) are transcripts which are co-expressed with their sense(S) counterparts without encoding detectable proteins or peptides. S/AS co-expression, no longer restricted to imprinted loci seen in chromosome inactivation, is a phylogenetically conserved phenomenon(Pontier & Gribnau, 2011). Suggested by large-scale RNA-Seq technologies and high-speed computational analyses, it is estimated that 26% of protein-coding sequences possess overlapping NATs which probably represent 29% of long non-coding RNA (lncRNA, >200 nucleotides in length) in humans (Necsulea et al., 2014; Ning et al., 2017). However, these data fail to reveal the presence of NATs in inducible genes. For example, *FCERIA*-AS reported in this study is not found in several databases examined probably due to expression of *FCERIA*-S is induced. By using IL-3 induction, we are able to identify an almost identical co-expression pattern between *FCERIA*-AS lncRNA and *FCERIA*-S.

We found that *FCERIA*-AS is fully overlapped with sense transcript. To our knowledge, this is the first time that fully overlapped AS is reported. The mechanism for expression of S/AS side by side is not well known. It seems that the putative promoter region of antisense direction of *FCERIA* are in relaxing state as in sense direction in cells being able to express this molecule. GATA-2 only plays weak promoting activities, indicating other initiation force of antisense transcription may exist. Bi-directional promoters probably rich in CpG-island may allow cognate NATs being produced with so called divergent head-to-head configuration (Balbin et al., 2015). It is hard to imagine this fashion can result generation of fully overlapped NATs. The other proposed model is that mobile genetic elements especially ancient transposon sequences like MIR and L2 which may be present upstream of antisense sequences, functioning as promoter to trigger the transcription of AS when chromosome structure of the locus relaxes (Conley, Miller, & Jordan, 2008; Zinad, Natasya, & Werner, 2017). Sequence blast analysis revealed that transposon MIR element is spread within this region (Figure7-supplementary1). Additionally, the transcription of FCERIA-AS is due to extension of transcription of neighboring gene located 3’ down stream of FCERIA. This possibility is excluded since RNA-Seq data from FcεRIα-expressing cells failed to collect any signals from the closest downstream neighbor gene Olfr1404 (Figure7-supplementary1).

Additionally, absence of *FCERIA*-AS in both of our gene manipulated mouse models may suggest that sequence-intrinsic mechanisms are critical to initiate or maintain the presence of antisense transcription. Indeed, the nascent transcript for antisense transcripts was also defective in Model II animals, indicating a requirement for sequence-dependent initiation of antisense transcription (data not shown). Conceivably, these unknown sequence-intrinsic mechanisms can become uncertainties during gene manipulation endeavor. For example, failure of expression of mouse siglec H was reported when DTR was inserted before 3’UTR of this gene (Takagi et al., 2011). Insertion of human dihydrofolate reductase (hDHFR) resistance cassette downstream of 3’UTR of sexual commitment gene GDV1 in malaria parasite disrupted the expression of GDV1 antisense RNA, which subsequently resulted dysregulated expression of sense GDV1 (Filarsky et al., 2018).

Based on a few examples of transcriptional interference of sense transcript by AS reported, AS can only regulate expression of sense cognate on the same chromosome, so called cis-regulation(Guttman & Rinn, 2012). In Model I reported in this study, loss of *FCER1A*-S and *FCER1A*-AS transcripts was found in homozygous mice whereas heterozygous mice still possessed S/AS transcripts expression with reduced levels (Fig 4c, 4e, 5a, 5b, 5c, and 5d). These findings strongly indicate expression of *FCER1A*-AS from sister chromosome is unable to promote inhibited transcription of sense transcripts on the other chromosome. These, therefore, provide another strong paradigm for regulation of S/AS via cis-acting manner which only affects gene expression monoallelically. Epigenetic-like mechanisms have been proposed in understanding the cis regulation of sense transcription by antisense (Zinad et al., 2017).

Both upregulation and downregulation are seen in terms of regulation of sense gene expression by its NATs. X chromosome inactivation regulated by XIST/TSIX offered the earliest and best studied example on repression of gene activity by NATs (Pontier & Gribnau, 2011). We found, however, a depressing effect played by *FCERIA*-AS on sense gene expression, since presence of AS is required for the induction its sense partner. Single strand RNA knockdown of *FCER1A*-AS via CasRx strategy resulted in reduction of FCER1A mRNA and protein. Similar positive regulatory effects have also been reported on tumor suppressor gene PDCD4 control of breast cancer progression and association of BACE1 with Alzheimer’s disease pathophysiology, in which mRNA stability is found to be promoted by their NATs (Faghihi et al., 2008; Jadaliha et al., 2018). In this investigation, we speculated that *FCER1A*-AS may promote FCER1A mRNA expression through transcriptional initiation.

FCER1A is a very peculiar gene for its existence, since it is not essential for survival and fertilization(Dombrowicz, Flamand, Brigman, Koller, & Kinet, 1993) with extremely limited expression spectrum on basophils and mast cells under steady state. Its existence seems only bring us IgE-dependent allergic reactions to expel unpleasant antigens in some individuals. Dysregulated allergic reactions result IgE-dependent allergic diseases which can be detrimental. Fcε RIα expression is promoted by relevant cytokines like IL-3 and allergic inflammation (Wang et al., 2021). The fine balance between good and ill outcomes of allergic reactions calls for stringent control of its expression. In addition to well-known trans regulatory molecules like GATA-2 and STAT-5(Li et al., 2015; Ohmori et al., 2015) which control Fcε RIα expression bi-allelically, our findings provide another layer of control on Fcε RIα expression monoallelically. The high effectiveness on downregulation of Fcε RIα expression shown here may suggest *FCER1A*-AS being another potential target for anti-allergic drugs.

## Materials and Methods

### Mice

Wild type C57BL/6 mice were purchased from SLAC Laboratory (Shanghai, China). All mice were housed and bred under specific pathogen free conditions in the Animal Center of Tongji University and used at 6–8 wk of age. All procedures performed on animals within this study were approved by the Committee on the Ethics of Animal Experiments of Tongji University (permit numbers: TJAA07920101)

FCER1A^dtr/dtr^ mice were generated on B6 background by Cyagen (Suzhou, China) using homologous recombination strategy depicted in Fig.3A. The linearized targeting vector containing a P2A-DTR-loxP-Neo^r^-loxP auto-deleter cassette was delivered to ES cells (C57BL/6) via electroporation. DTR-containing cassette was placed into *FCERIA* allele by homologous recombination with homology arms flanking 4.8-kilobase upstream of exon 3 and 2.8-kb downstream of exon 5 respectively, allowing P2A-DTR being inserted in-frame before stop codon. The primers for genotyping were listed in table1.

FCER1A^dtrGFP/dtrGFP^ mice were generated on B6 background by Biocytogen (Beijing, China) using Cas9/sgRNA strategy depicted in Fig.3B. The targeting construct, assembled by PCR and cloning, consists of two homology arms and sequence of DTR-P2A-EGFP with sequences of 5’UTR and 3’UTR being remained intact. One arm is 1.5-kb fragment spanning upstream of exon 1 up to start codon containing whole 5’UTR, and another arm is 1.5-kb fragment downstream of stop codon containing whole 3’UTR. Two sgRNAs were designed to target at regions proximal to upper and lower homology arms respectively as illustrated in Figure3. sgRNAs and Cas9-expressing vectors along with targeting constructs were delivered to fertilized zygotes through microinjection. The primers for genotyping were listed in table S1. Homozygous mice of FCER1A^dtrGFP/dtrGFP^ is used as knock out of FCER1Ain this study. Mice of both lines are fertile and healthy.

### Infection

Gender- and age-matched mice were infected percutaneously with 30 cercariae of *S. japonica*, which were shed from infected Oncomelania hupensis snails provided by National Institute of Parasitic Diseases at Shanghai, China.

### Cell culturing

Bone marrow (BM) cells were flushed from femurs and tibiae of 6-10 weeks age of mice of wild type, FCER1A^dtr/dtr^, or FCER1A^dtrGFP/dtrGFP^. After lysis of red blood cells, isolated cells were cultured in RPMI 1640 medium (GIBCO, Invitrogen) containing 10% heat-inactivated FBS (GIBCO, Invitrogen), 50 μM 2-mercaptoethanol (GIBCO) at 1×10^6^ /ml. 10 ng/ml IL-3 was added for induction of Fcε RIα-expressing cells, or 10ng/ml GM-CSF was added for induction of non Fcε RIα-expressing cells with medium being replaced every 2 days. The cells usually were harvested 6 days after culture.

Mast cell line MC/9 was purchased from ATCC(CRL-8306) cultured in Dulbecco’s Modified Eagle’s Medium(GIBCO, Invitrogen) adjusted to contain 4 mM L-glutamine, 4.5 g/L glucose,1.5 g/L sodium bicarbonate and supplemented with 2 mM L-glutamax(GIBCO, Invitrogen), 50 μM 2-mercaptoethanol, 10% concanavalin A(Sigma-Aldrich) stimulated-splenocytes supernatant, 10 ng/ml IL-3 (Peprotech), and 10% fetal bovine serum(GIBCO, Invitrogen).

### Generation of plasmids

The vectors for expression of CasRx and sgRNA were kindly provided by Dr. Hui Yang (Institute of Neuroscience, Chinese Academy of Sciences, Shanghai, China). To construct vectors expressing sgRNA sequences of interest, backbone vectors of U6-DR-sgRNA-DR-EF1a-mCherry-SV40 were digested with NdeI and AsiSI restriction enzymes (NEB) first, followed by ligation with inserted sequences containing various sgRNA (Table S2) which were synthesized by Sangon Biotech Company (Shanghai). MC/9 mast cells were co-transfected by electroporation with 7ug of CasRx vector and 11ug of sgRNA vector. 5×10^6^ MC/9 cells were suspended in 0.5 ml of optiMEM (GIBCO) and kept at room temperature for 10min before electroporation was performed as stated above.

Putative regions for mouse *FCER1A*-S promoter and mouse *FCER1A*-AS promoter were synthesized by Sangon Biotech Company (Shanghai) and inserted into the multicloning sites of pGL-4.17 Basic (Promega) to generate reporter plasmids for detection of putative promoter activities. GATA-2-expressing vector was constructed by cloning synthesized mouse GATA-2 sequences into eukaryotic expression vector pcDNA3.1 (+) (Invitrogen) with restriction enzymes of NheI and EcoRI (NEB). HEK293T cells were transfected with 3ug pGL-4.17 reporter plasmid, 3ug Gata-2 expression plasmid, and 375ng hRluc (Promega) by Lipofectamine 3000 (Thermofisher). Luciferase activity was determined using dual-luciferase assay kit (Promega) as described (Maeda et al., 2006).

### Transfection of cells

5 × 10^6^ MC/9 cells were suspended in 0.5 ml of optiMEM (GIBCO) and kept at room temperature for 10 min. Cells were then electroporated using Gene Pulser II® (Bio-Rad) with pulse conditions of 800 µF and 280 V. Cells were kept an additional 10 min at room temperature before being placed back to 37°C incubator in MC/9 culturing medium. All detection was made 48 hours after transfection.

### Strand-specific reverse transcription and quantitative real-time PCR

Total RNA was prepared by using RNAiso Plus (Takara). 1 μg of total RNA was used to synthesize cDNA with PrimeScript RT reagent kit (RR047A, Takara) according to the manufacturer’s instructions. Briefly, RNA was treated with gDNA Eraser (Takara) to remove genomic DNA first before1ug of total RNA was reverse transcribed for 15 or 20 min either with random hexamers or with strand-specific primers of *FCREIA* (Table S1). PCR was subsequently performed to amplify cDNAs at 35 cycles with various annealing temperatures. PCR products were visualized on agarose gels by GoldView. Quantitative real-time PCR was performed using SYBR Premix EX Taq (Tli RNaseH Plus) (Takara) in a QuantStudio3 thermal cycler (Applied Biosystems, Foster City, CA). The qPCR values for GATA-2 and CD200R3 were all normalized first by GAPDH. The specific primers employed were shown in Table S1

### Strand-specific RNA sequencing

RNA from either IL-3 induced bone marrow cells or MC/9 cells was isolated as stated. 5 μg RNA was used as input material for strand-specific RNA-Seq. Firstly, ribosomal RNA was removed by Epicentre Ribo-zero™ rRNA Removal Kit (Epicentre) and rRNA-free residue was obtained by ethanol precipitation. Sequencing libraries were subsequently generated by NEBNext® Ultra™ Directional RNA Library Prep Kit for Illumina® (NEB). Briefly, after RNA fragmentation, rRNA-free RNA was subjected to first-strand cDNA using random hexamer primer and M-MuLV Reverse Transcriptase. Second-strand cDNA synthesis was performed using DNA polymerase I and RNase H with dTTP being replaced by dUTP. After end repair, polyA tailing, adaptor sequence addition and U excision, the strand specific libraries were sequenced by Novogene Bioinformatics Institute (Beijing, China) on an Illumina Hiseq 4000 platform and 12G of paired-end 150 bp reads were generated.

### RNA-Seq Data Analysis

Sequence data from MC/9 libraries was aligned to mouse genome build mm10 using Hisat2 software 2.2.1 (Kim, Paggi, Park, Bennett, & Salzberg, 2019; Pertea, Kim, Pertea, Leek, & Salzberg, 2016). SAMtools 1.14(http://www.htslib.org/) was applied to detect and analyze regions of signal on each strand of *FCERIA* location. Bam2wig.py converts all types of RNA-seq data from BAM format into wiggle format. Data visualization was aided by IGV software based on information of reads depth.

### ChIP-seq and ATAC-seq data analysis

To identify the epigenetic profile of FCER1A locus, we queried NCBI Gene Expression Omnibus (GEO) datasets. For data within NCBI (GSE145542 and GSE145544), the data in TDF format file was retrieved from GEO. The analyzed sequence data was visualized with IGV software

### Rapid amplification of cDNA ends (RACE)

5’ and 3’ RACE was performed with SMARTer RACE 5’/3’ Kit (Takara) following the manufacturer’s instructions. In brief, total RNA was extracted from IL-3 induced bone marrow cells or MC/9 cells. First-strand cDNA was synthesized with gene specific primers as shown in Table S1. Touchdown PCR was then performed to amplify different transcripts. Adding poly (A) tails to RNA (Takara) following the manufacturer’s instructions. PCR products were gel extracted for Sanger sequencing.

### Capture of Nascent RNAs

To capture nascent RNAs, Click-it Nascent RNA Capture Kit (Life Technologies) was used as manufacturer’s instructions. Briefly, on the 6th day of IL-3 induced bone marrow cell culturing, 0.5 mM 5-ethynyl uridine (EU) was added into culture medium. 30 minutes after incorporation, total RNA was prepared with RNAiso Plus (Takara). 1μg EU-labeled RNA was biotinylated with 0.5 mM biotin azide in Click-iT reaction buffer for 30min. The biotinylated RNAs were then precipitated with ethanol and resuspended in DNase/RNase-free distilled water. The biotinylated RNAs were captured by streptavidin T1 magnetic beads (Dynabeads MyOne) in Click-iT RNA binding buffer at 68-70°C for 5 min, followed by incubation at room temperature for 30min while gently vortexing. The beads were immobilized using the DynaMag-2 magnet and then were washed with 5 × Click-iT wash buffer 1, followed by 5 × wash buffer 2, and resuspended in a wash buffer 2. The strand-specific reverse transcription for captured nascent RNA transcripts and subsequent quantitative real-time PCR were performed as manufacturer’s instructions

### Flow cytometry analysis of surface molecules and intracellular cytokines

FACS staining for surface molecules and intracellular cytokines for cultured cells or splenocytes were performed as described previously (Ma et al., 2015). For peripheral blood cells, orbit blood was collected from isoflurane anesthetized mice. Blood drawing apparatus were coated with 0.1 M EDTA to prevent clotting. The blood samples were then subjected to fluorescence-labeled antibody staining before RBC was dissolved by lysing solution (LSB01, MultiSciences Biotech, Hangzhou, China). For various fluorescence-labeled antibodies and reagents for live cell detection were listed in Table S3. Final fluorescence detection was made by FACSVerse (BD Biosciences) or CytoFLEX LX (Beckman Coulter). Flow cytometry data analysis was analyzed by FlowJo software (FlowJo) or CytExpert (Beckman Coulter).

### Passive cutaneous anaphylaxis (PCA)

PCA was performed as reported (Cruse et al., 2016; Dombrowicz et al., 1993). Animals were primed intradermally in both sides of ear pinna with 100μg of murine anti-DNP IgE (Sigma-Aldrich) dissolved in 20 ul PBS. 20hrs later, 500μg DNP-BSA (Sigma-Aldrich) in 100μl PBS containing 1% Evans blue dye was injected through tail vein. Animals were euthanized 1.5hrs after DNP-BSA challenge for dye extravasation assessment at primed ears. Both ears were removed and minced and placed into glass vials. The dye contained were extracted by dissolving the shredded tissues in 0.5mL dimethylformamide (DMF). Vials were then shaken at 500 rpm of 55°C for 3 to 5 hrs. Optical density of Evan blue in each ear was measured at 650 nm to reflect the degree of extravasation.

### Statistics

Graphs were generated by GraphPad Prism 7 (GraphPad Software). Statistics were performed as illustrated in relevant figure legends with significance as *p<0.05, **p<0.01, ***p<0.001 and ****p<0.0001.

## Acknowledgments

The authors thank Dr. Zhang qingfeng for helpful and insightful input on the role of antisense regulation.

This work was funded the National Natural Science Foundation by China (grants 81772215 and 81572021 to X.-P. Chen).

## Author contributions

R.Y. Tang and X.-P. Chen conceptualized and designed the study. R.Y. Tang and X.-P. Chen wrote the paper. R.Y. Tang conducted a majority of the experiments. L. Yin analyzed the FACS data. L. Yao genotyped animals.

The authors declare no competing financial interests.

## Figure legend

**Figure1-Supplementary1:**
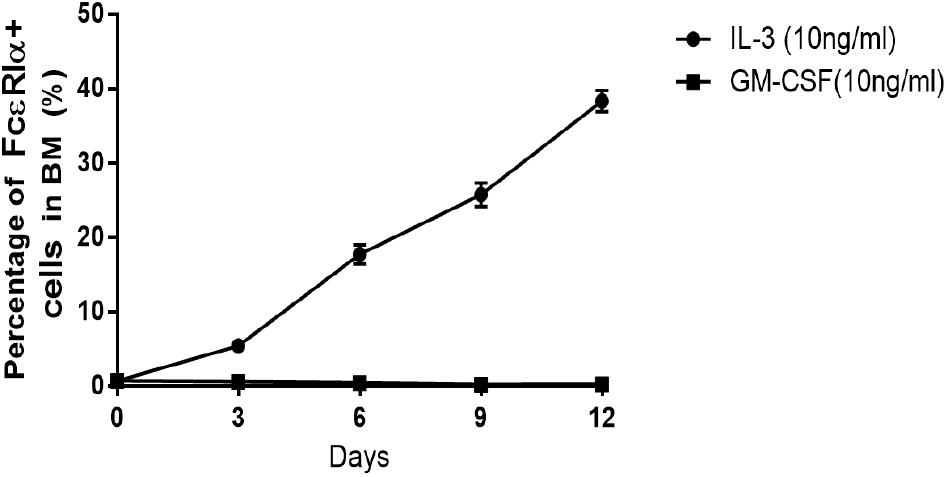
Time course of FcεRIα-positive cells in IL-3 (circle) - or GM-CSF (square)-treated bone marrow cells. Combined data from 4 independent mice show the percentage of FcεRIα-positive cells by FACS staining at different time point in relation to that at time zero.

**Figure1-Supplementary2:**
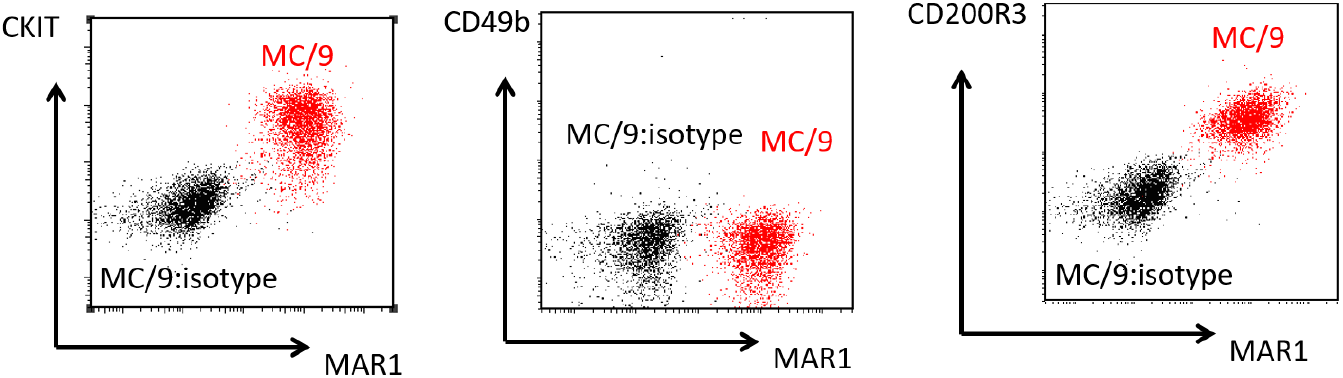
Cell surface expression of c-kit, FcεRIα and CD200R3 on MC/9.

**Figure1-Supplementary3:**
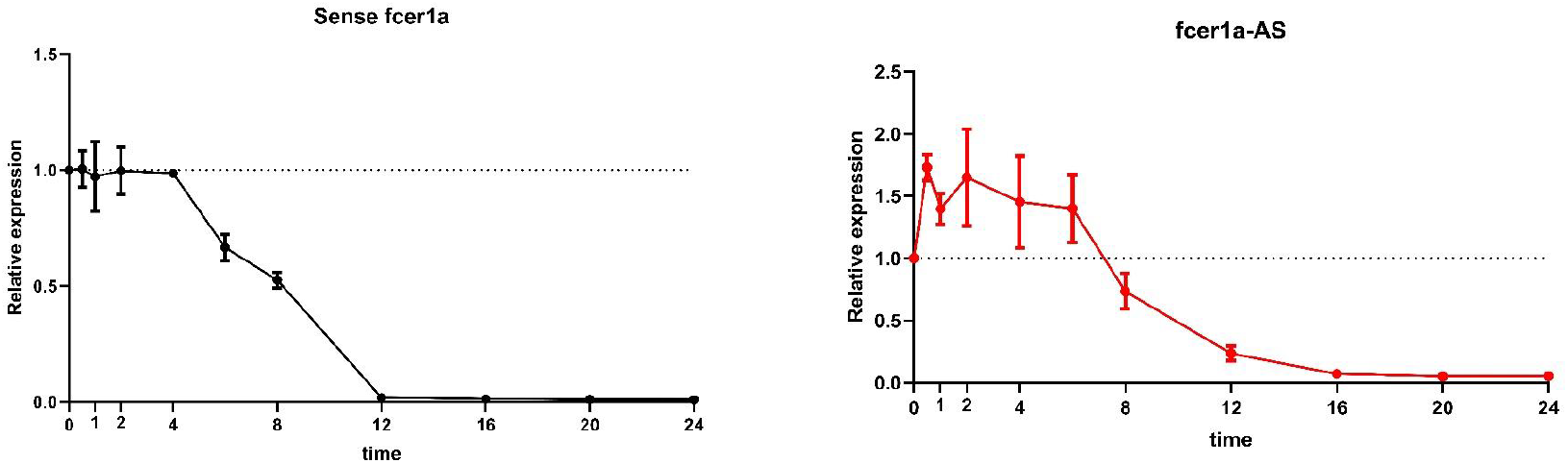
RNA stability of FCER1A-S and FCER1A-AS transcripts at different times points after actinomycin-D treatment. Amount of FCER1A-S and FCER1A-AS transcripts over time (at 0, 0.5, 1, 2, 4, 8, 12, 16, 20 and 24 h) after actinomycin-D treatment was measured by RT-PCR in MC/9 cells. Shown are expression levels at each tome point relative to time 0. Data are mean ± SD of three different experiments.

**Figure1-Supplementary4:**
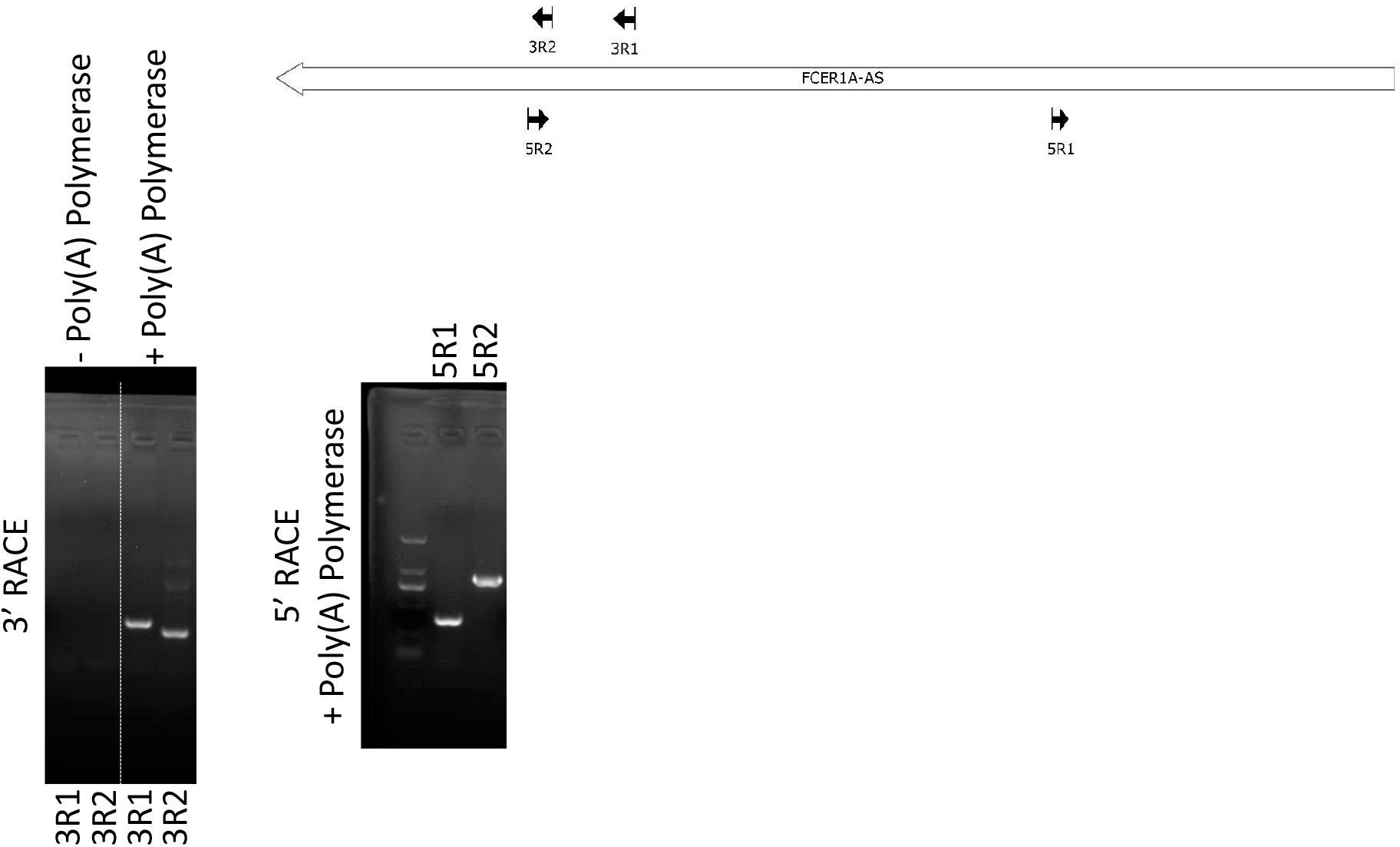
Identification of the transcriptional starting site (TSS) and ending site of FCER1A-AS in MC/9 cell lines (A) 3’RACE fails unless poly (A) tails is added at 3’end of antisense RNA. Transcriptional ending site identified with 3’RACE assay in PolyA-added RNA. TSS identified with 5’RACE assay in PolyA-added RNA. Primers 3R1/3R2 and 5R1/5R2 indicated the primers used in RACE assay.

**Figure1-Supplementary5:**
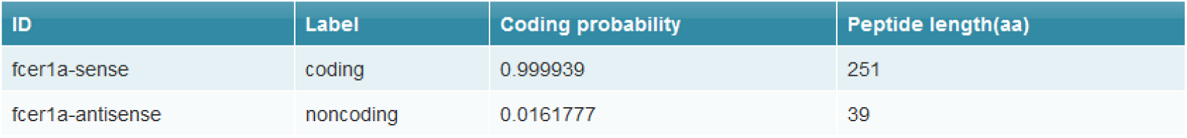
Bioinformatics analysis suggested that FCER1A-AS had no coding capability (http://cpc2.gao-lab.org/)

**Figure2-Supplementary1 :**
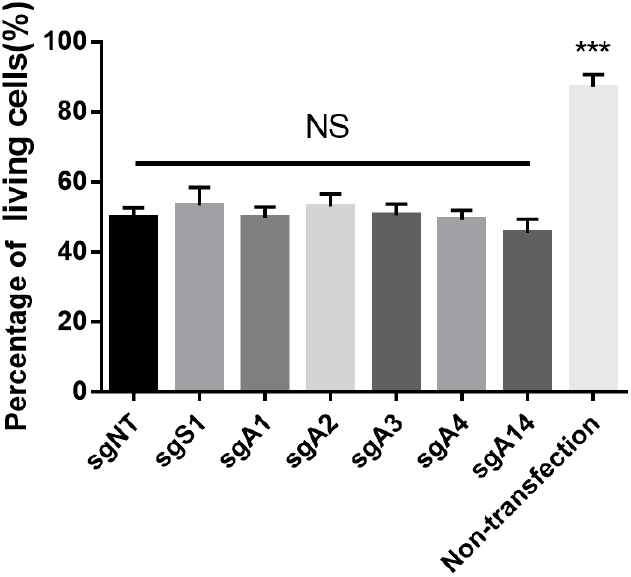
Transfection of CasRX and sgRNA sgNT∼sgA14 by electroporation did not affect viability differently. n=3 independent experiments. NS: no significant differences

**Figure2-Supplementary2:**
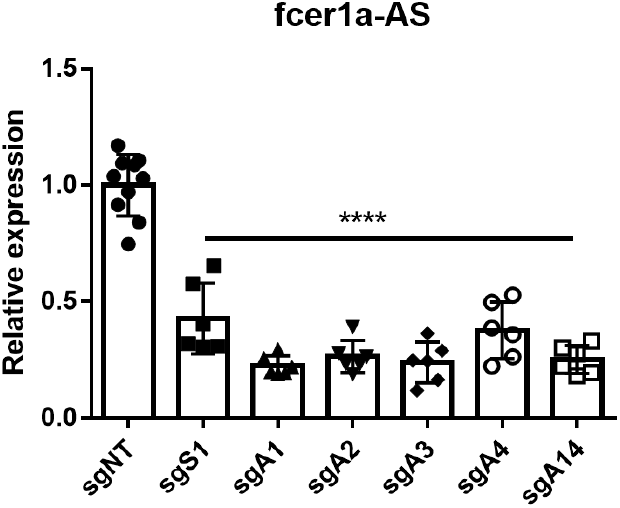
FCER1A-AS expression is successfully knocked down by FCER1A-AS-sgRNA (sgA) in MC/9 cells. Strand-specific quantitative RT-PCR of FCER1A-AS expression following FCER1A-AS-sgRNA (sgA) or FCER1A-S-sgRNAs (sgS) transfection. Relative expression indicates the ratio relative to control sgNT. Data are mean ± SD of six different experiments. ****P < 0.0001.

**Figure2-Supplementary3 :**
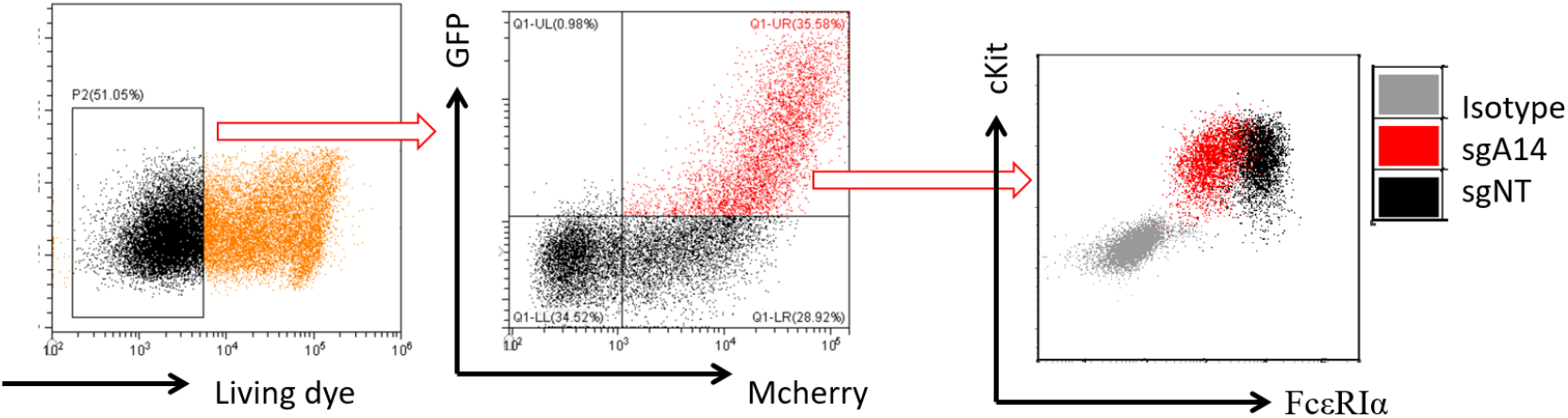
Gating Strategy for CasRX/sgRNA co-transfected cells.The GFP-positive (CasRX transfected) and Mcherry-positive (sgRNA transfected) population was selected on viable cells. In the case shown here, the cell surface expression of FcεRIα was markedly reduced by sgRNA A14 targeting onto FCER1A-AS comparing to non-targeting sgRNA NT control.

**Figure5-Supplementary1:**
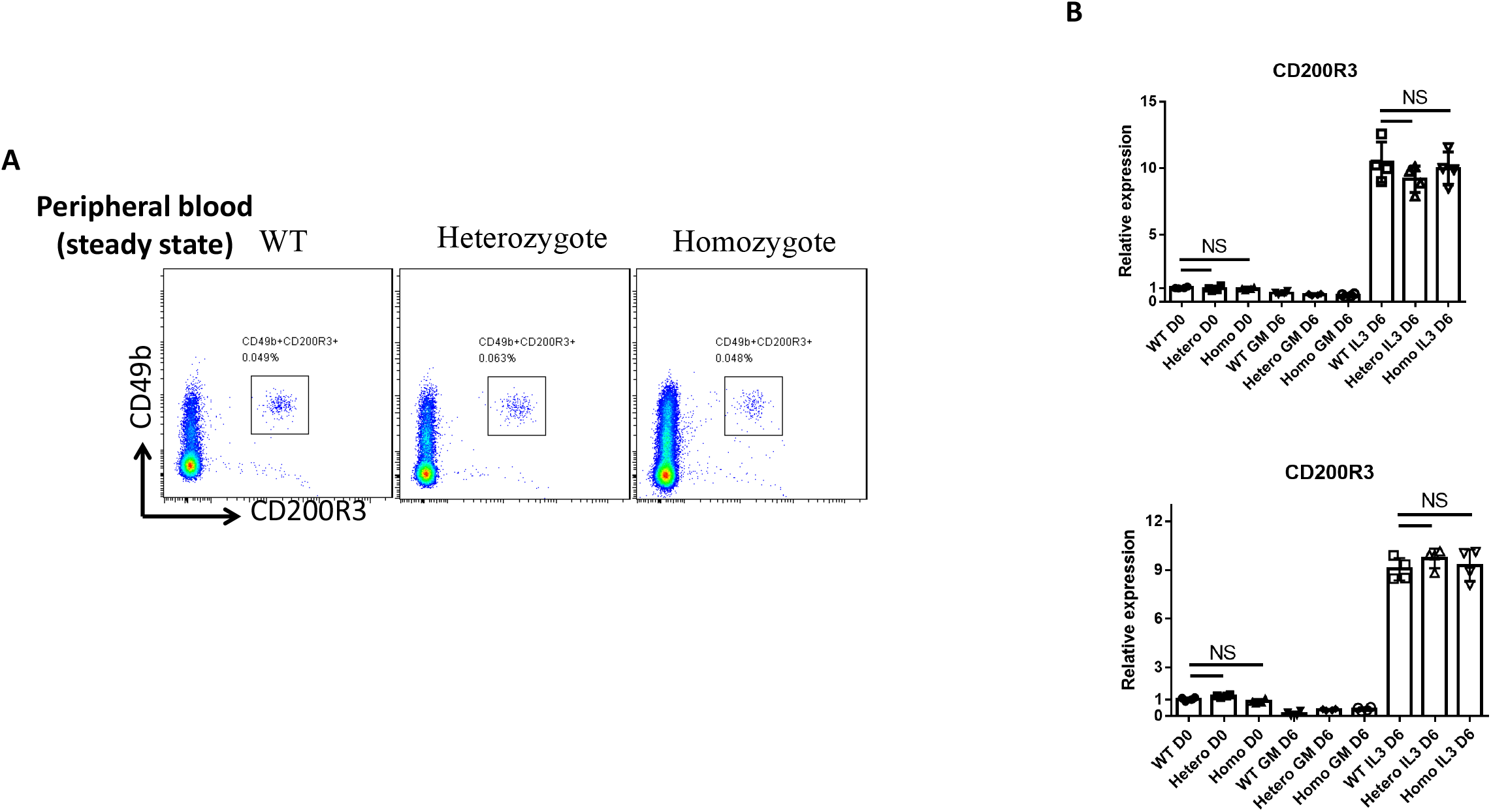
Basophils are not affected in Model I and Model II. (A) Basophils in peripheral blood were measured by expression of CD49b and CD200R3 in mice of WT, heterozygous Model I and homozygous Model I. (B) Basophils differentiation was examined from 6 days of IL-3-treated bone marrow cells in WT, heterozygous and homozygous mice of both Model I (upper) and Model II (lower) by qRT-PCR of CD200R3. RNA from each group was reverse transcribed by random hexamers. CD200R3 expression levels shown are relative to WT bone marrow cells at day 0. Data are the mean ± SD of four separate animals from each model.

**Figure6-Supplementary1:**
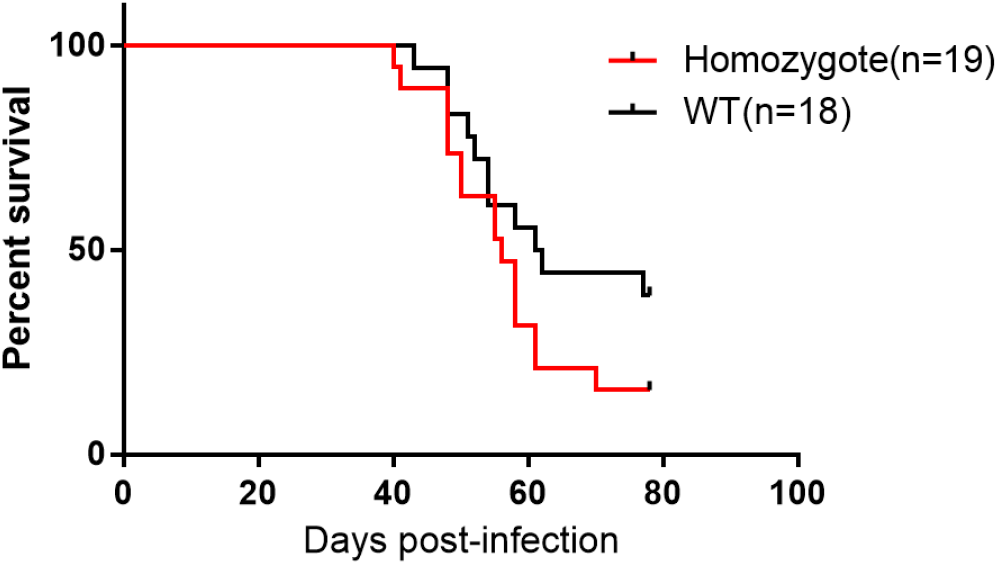
Survival kinetics of Schistosome japonica (Sj)-infected WT, FCER1A^dtr/dtr^ (Model I) mice. Combined data from three independent experiments with WT, n=18; FCER1A^dtr/dtr^ n=19 by prism 7.

**Figure7-Supplementary1:**
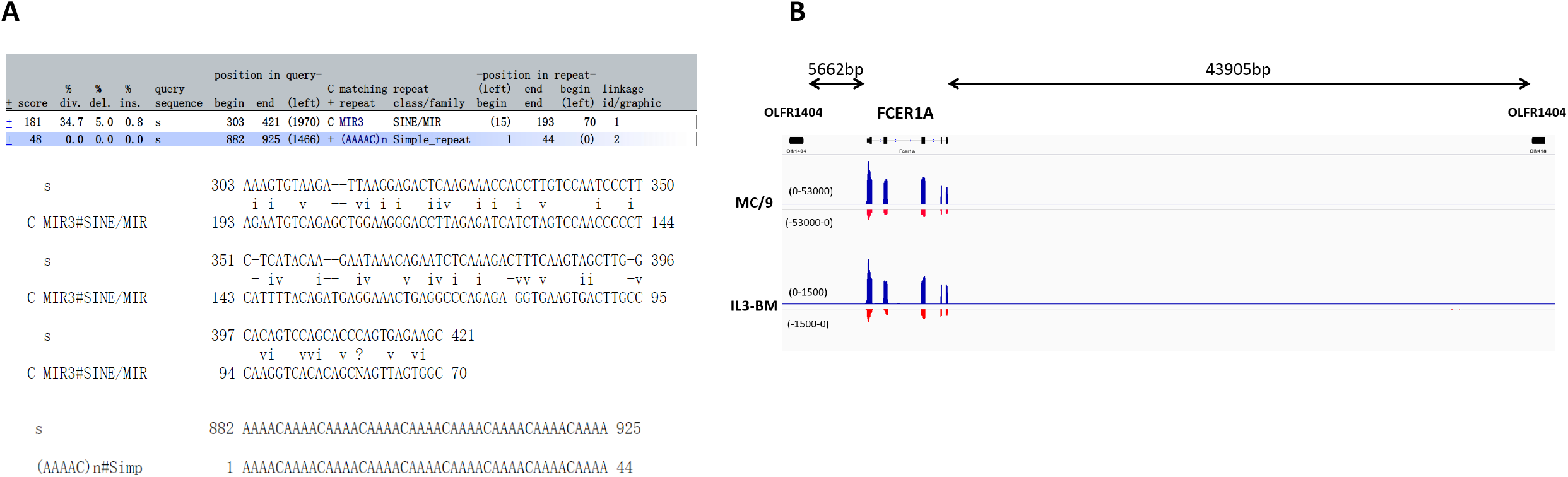
Putative transcription initiation analysis for FACERIA-AS. (A) Transposon MIR element and simple repeat sequence is spread within this promoter region of FCER1A-AS. (B)The transcription of FCERIA-AS is not due to extension of sense gene transcription of neighboring gene located 3’ down stream of FCERIA.

## Extend data

**Table S1:**
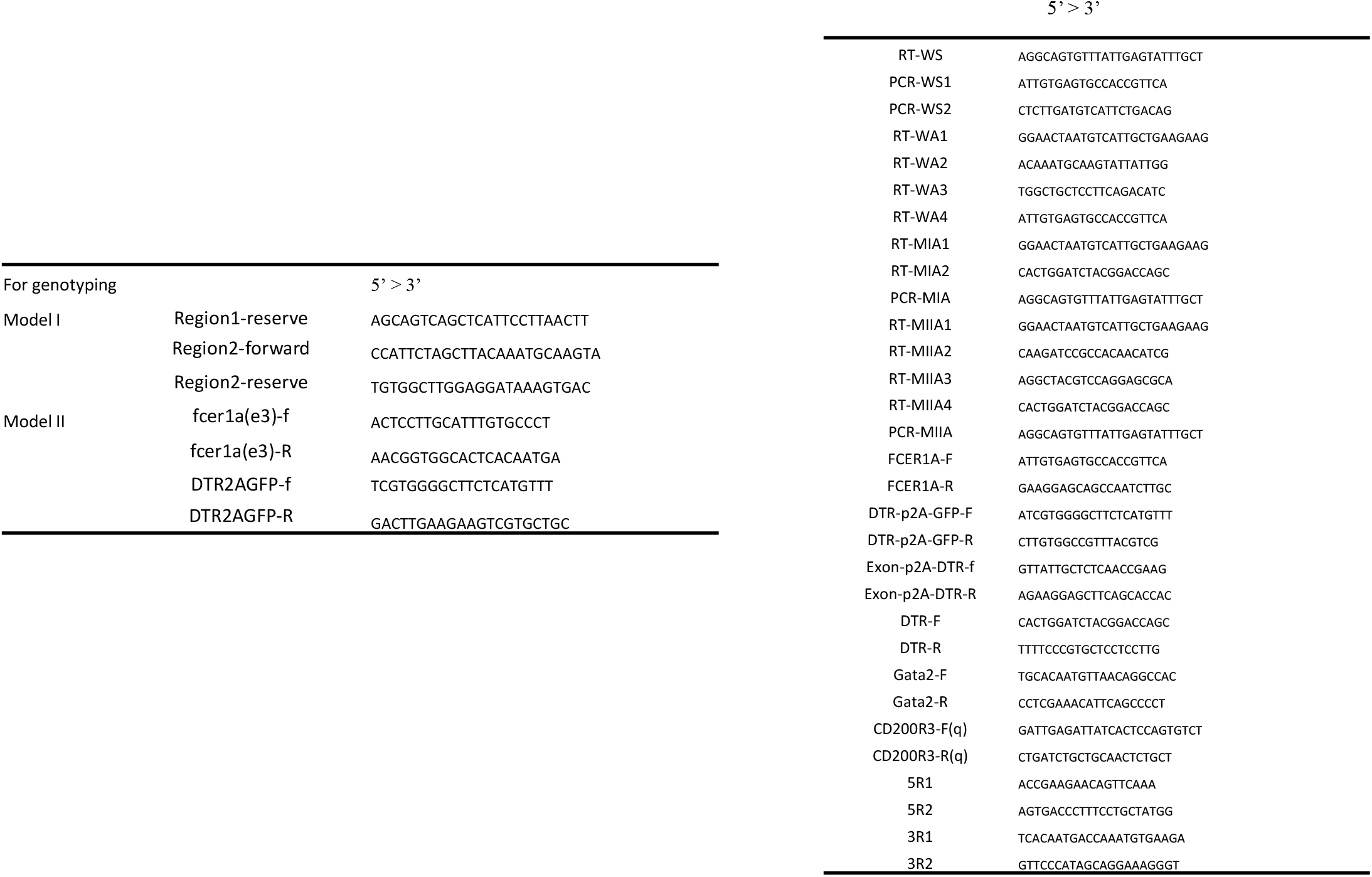
Primers used in this study

**Table S2:**
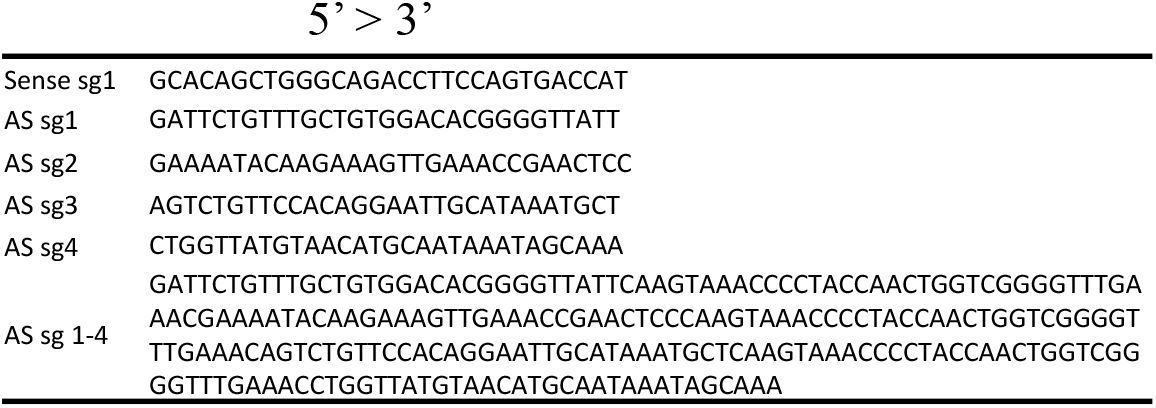
sgRNA sequence

**Table S3:** Fluorescence-labeled antibodies and reagents for live cell detection

|  |
| --- |
| APC-labeled anti-mouse FcεRIa (MAR-1; BioLegend) |
| PE-cy7-labeled anti-mouse CD49b (DX5; eBioscience) |
| PE-labeled anti-mouse Ckit(Ba13, BD Pharmingen) |
| PE-labeled anti-mouse CD200R3 (ACK2, BD Pharmingen) |
| FITC-labeled anti-mouse CD3ε (145-2C11; BioLegend) |
| Percp-Cy5.5–labeled anti-mouse B220 (RA3-6B2; BioLegend) |
| Fixable viability dye eFluor 780 (eBioscience) |

